# Hormonal coordination of peripheral motor output and corollary discharge in a communication system

**DOI:** 10.1101/2023.04.10.536282

**Authors:** Matasaburo Fukutomi, Bruce A. Carlson

**Author notes:** Corresponding Author Bruce A. Carlson. **Author Contributions** M.F. and B.A.C. designed research; M.F. performed research; M.F. analyzed data; and M.F. and B.A.C. wrote the paper. **Competing Interest Statement** The authors declare no competing interest.

## Abstract

Steroid hormones remodel neural networks to induce developmental or seasonal changes in animal behavior, but little is known about hormonal modulation of sensorimotor integration. Here, we investigate hormonal effects on a predictive motor signal, termed corollary discharge, that modulates sensory processing in weakly electric mormyrid fish. In the electrosensory pathway mediating communication behavior, inhibition activated by a corollary discharge precisely blocks sensory responses to self-generated electric pulses, allowing the downstream circuit to selectively analyze communication signals from nearby fish. These electric pulses are elongated by increasing testosterone levels in males during the breeding season. Using systematic testosterone treatment, we induced electric-pulse elongation in fish and found that the timing of electroreceptor spiking responses to self-generated pulses (reafference) was delayed as electric pulse duration increased. Recording evoked potentials from a midbrain electrosensory nucleus revealed that the timing of corollary discharge inhibition was delayed and elongated by testosterone. Further, this shift in corollary discharge timing was precisely matched to the shift in timing of the reafferent spikes. We then asked whether the shift in inhibition timing was caused by direct action of testosterone on the corollary discharge circuit or plasticity of the circuit through altered sensory feedback. We surgically silenced the electric organs of fish and found similar hormonal modulation of corollary discharge timing between intact and silent fish, suggesting that sensory feedback was not required for this shift. These results demonstrate that testosterone directly and independently modulates peripheral motor output and a predictive motor signal in a coordinated manner.

**Significance:** Self-other discrimination is essential for animals. Internal predictive motor signals, or corollary discharge, provide motor information to sensory areas so that animals can perceive self- and other-generated stimuli differently. As behavior and associated sensory feedback change with development, corollary discharge must adjust accordingly. Using weakly electric mormyrid fish, we show that the steroid hormone testosterone alters electric signaling behavior and the resulting sensory feedback, as well as the timing of corollary discharge, to precisely match the altered sensory feedback. We also found that the altered sensory feedback itself is not necessary to drive this corollary discharge modulation. Our findings demonstrate that testosterone directly and independently regulates peripheral motor output and corollary discharge in a coordinated manner.

## Introduction

Steroid hormones underlie seasonal or developmental changes in animal behavior, such as the seasonal songs of birds and the lowering of the voice in humans through secondary sexual characteristics. Such behavioral shifts are based on direct effects of hormones at multiple sites including peripheral effectors (1–4), central motor circuits (5–7), and sensory systems (8–10). Because these elements are closely interrelated, hormonal changes in behavior should require coordinated changes in sensorimotor integration. However, little is known about the hormonal control of sensorimotor integration. Here, we investigate hormonal effects on a corollary discharge that provides motor information to modulate central sensory processing in weakly electric mormyrid fish. In this system, mechanisms of corollary discharge and hormonal effects on motor output are well understood.

Mormyrid fish generate electric pulses by discharging an electric organ in their tail (11). These pulses are used for active electrolocation (12) and communication (13). The waveform of the electric organ discharge (EOD) is stereotyped to represent the sender’s identity, such as species, sex, and social status (1, 14–16), while the interval between EODs can be flexibly varied to communicate behavioral states in the moment (Fig. 1A) (17–19). Mormyrids have a dedicated sensory pathway for processing electric communication signals (20–22), in which a corollary discharge plays an essential role (23, 24). In this pathway, the primary sensory center (the nucleus of the electrosensory lateral line lobe [nELL]) receives two types of inputs: excitation from sensory afferents of electroreceptors (Knollenorgan [KO]) distributed throughout the surface of the skin, and inhibition originating from the EOD command nucleus (CN) in the medulla through a corollary discharge pathway (Fig. 1B) (21, 23–25). Both self- and other-generated EODs stimulate KOs, but this internal inhibitory signal precisely blocks sensory responses to self-generated EODs, allowing the downstream pathway to selectively process sensory information from the EODs emitted by nearby fish (Fig. 1B).

**Fig. 1.**
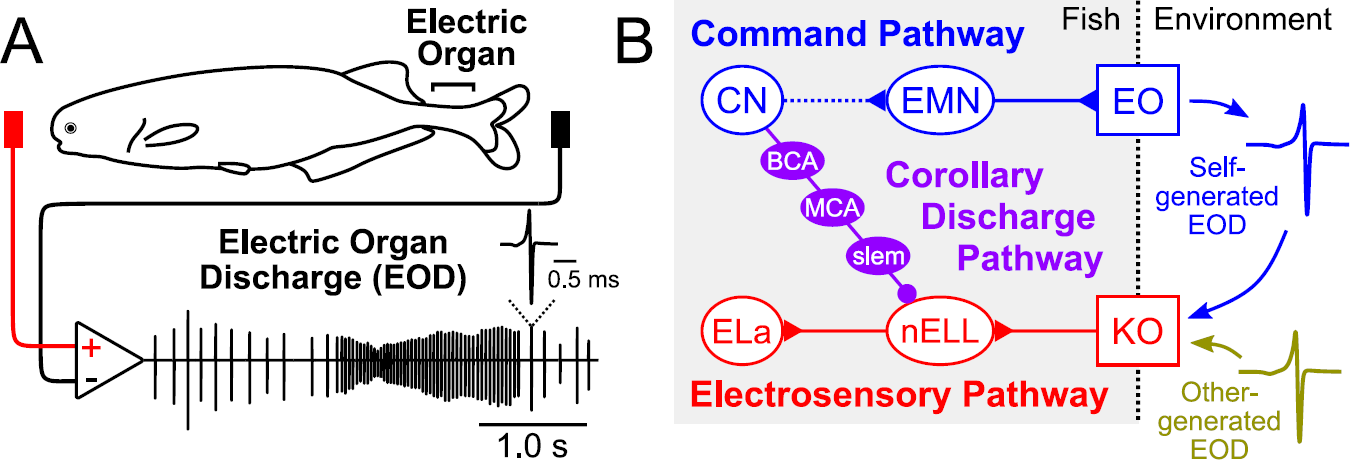
Electric signaling behavior and the sensorimotor circuit of mormyrid fish. (A) Electrical recording from a freely swimming mormyrid, *Brienomyrus brachyistius*. Electric signaling consists of a fixed waveform and variable inter-pulse intervals of electric organ discharge (EOD). The changes in EOD amplitude are due to movement of the fish relative to the recording electrode, not to changes in EOD amplitude emitted by the fish. If the recording electrode (red) is placed on the head side and the reference electrode (black) on the tail side, then a head-positive waveform will be recorded. (B) Circuit diagram showing electromotor command (blue), Knollenorgan (KO) electrosensory (red), and corollary discharge (purple) pathways. The command nucleus (CN) drives the electric organ (EO) to generate each EOD via the medullary relay nucleus (MRN, not shown) and spinal electromotor neurons (EMN). KO electroreceptors respond to both self- and other-generated EODs and send time-locked spikes to the first center (the nucleus of the electrosensory lateral line lobe [nELL]) via primary afferents. The CN sends corollary discharge inhibition to the nELL, via the bulbar command-associated nucleus (BCA), the mesencephalic command-associated nucleus (MCA), and the sublemniscal nucleus (slem), which blocks sensory responses to self-generated EODs. This circuit allows the nELL neurons to send filtered sensory information about EODs generated by other fish to the anterior exterolateral nucleus (ELa).

This communication system is sensitive to the steroid hormone testosterone (26). Exogenous administration of testosterone increases EOD duration in juveniles, females, and non-reproductive males, mimicking the sexual differentiation of mature males that occurs during the breeding season (1). In this case, testosterone directly affects the biophysical properties of the electrocytes in the electric organ that determine the EOD waveform (11, 27–29). Testosterone also induces a downward shift in the sensory tuning of the KOs to match the spectral content of the altered EOD, but this hormonal effect is indirect and depends on sensory feedback (30). If the sensory feedback (or reafferent input) changes, then the timing of corollary discharge inhibition would also need to change in concert to continue filtering out responses to self-generated EODs. Indeed, a previous study comparing different species of mormyrids showed that fishes with long EODs have delayed corollary discharge inhibition compared to those with short EODs (31). This difference in the timing of inhibition optimally matches the timing of reafferent input from KOs, which differs between species (31). Furthermore, this trend was observed for individual differences in EOD duration within one species (31). The present study aims to test whether the timing of this corollary discharge inhibition is altered by testosterone to match the shifted reafferent input, and the role of sensory feedback in mediating this change.

## Results

### Testosterone elongates EOD duration

We recorded individual EODs of freely swimming fish from the start of treatment to 13 days after treatment (Fig. 2A). Testosterone treatment increased EOD duration while the vehicle treatment had no effect (Fig. 2B; p = 0.0008 for treatment, p < 0.0001 for days after treatment, p < 0.0001 for the interaction, linear mixed model [LMM]). Accordingly, peak power frequencies of the EODs were lowered by the testosterone treatment (Fig. 2C; p = 0.0020 for treatment, p < 0.0001 for days after treatment, p < 0.0001 for the interaction, LMM). These results were consistent with previous studies (1, 28, 29). We also confirmed that testosterone treatment increased the delay to EOD peak 1 (Fig. 2D; p = 0.0009 for treatment, p < 0.0001 for days after treatment, p < 0.0001 for the interaction, LMM), which is strongly correlated with KO receptor spike timing in response to self-generated EODs (31).

**Fig. 2.**
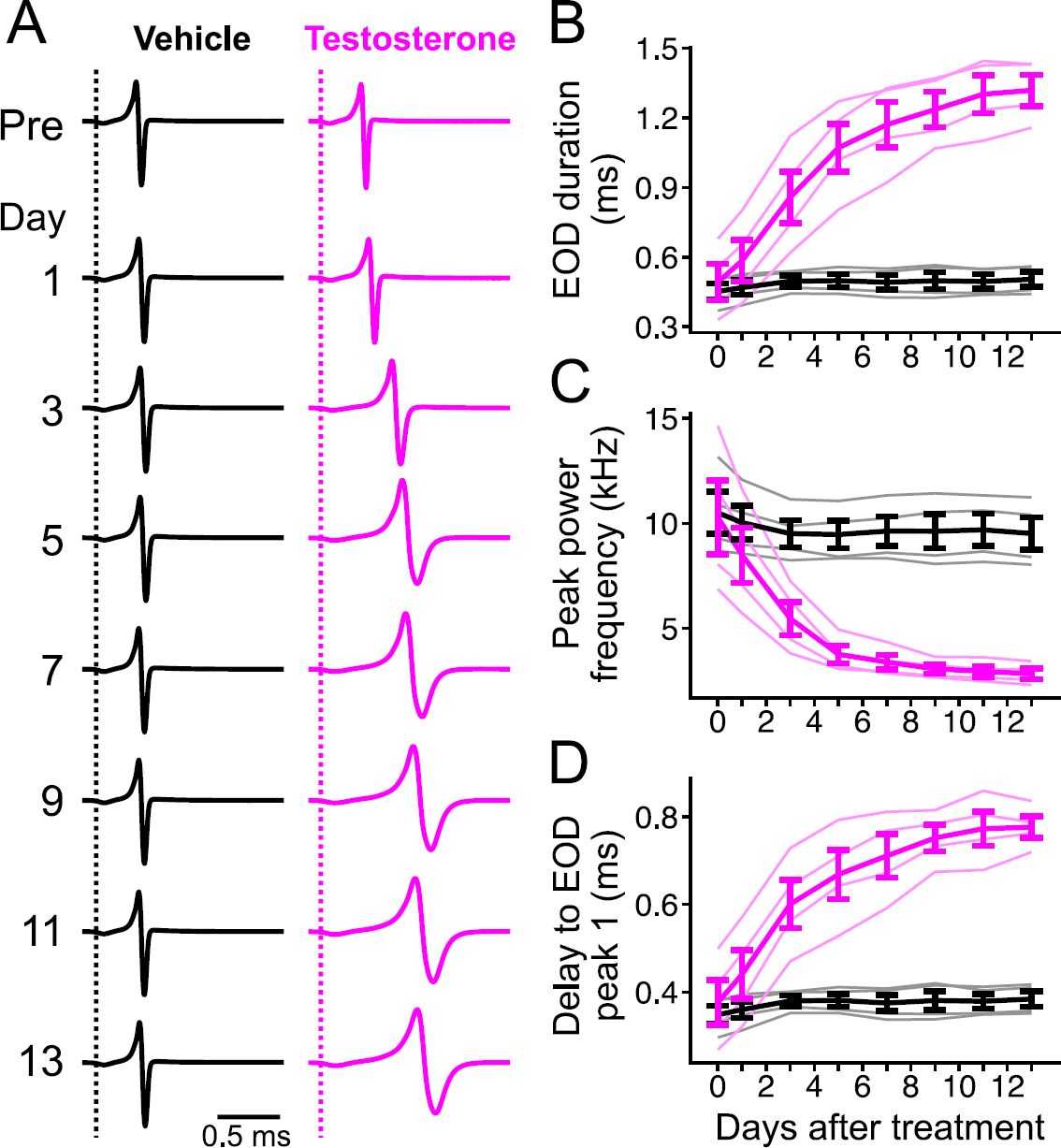
EOD duration is elongated by testosterone treatment. (A) Daily changes in EOD waveform in response to vehicle (black) and testosterone (magenta) treatment. Dotted line indicates EOD onset. (B–D) Daily changes in EOD duration (B), peak power frequency (C), and delay to EOD peak 1 (D). Each light-color line indicates individual fish and dark-color line indicates the average. Error bars indicate SEM.

### EOD elongation shifts KO spike timing

We recorded spiking activity from KOs of treated fish in response to electrosensory stimulation that mimics a self-generated EOD (Fig. 3A; see Materials and Methods). A representative KO produced a single spike with short delay following peak 1 of the EOD and was found to shift its spike timing along with testosterone-induced EOD elongation (Fig. 3B). Across KOs, we found that testosterone shifted the peak response latency (Fig. 3C; p < 0.0001 for treatment, p < 0.0001 for days after treatment, p < 0.0001 for the interaction, LMM). A linear regression comparing KO response latency with the delay to EOD peak 1 revealed a strong correlation, with a slope of 1.11 and intercept of 0.16 (Fig. 3D; p < 0.0001 for slope, p < 0.0001 for intercept, LMM). We also measured the correlation between KO peak latency and delay to EOD peak 1 and found a slope of 0.11, which was significantly larger than 0 (Fig. 3E; p = 0.033 for slope, LMM), indicating that KO spike latency relative to EOD peak 1 also slightly increased as the EOD elongated.

**Fig. 3.**
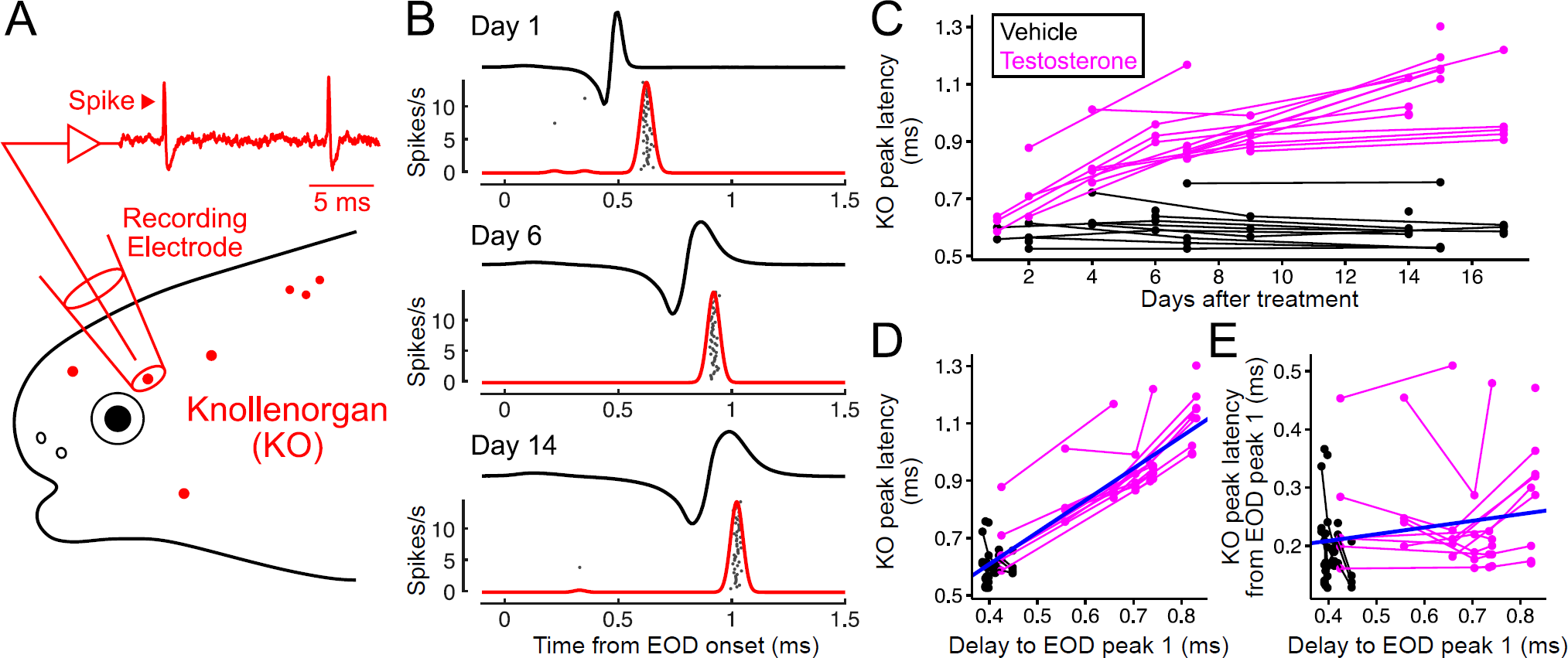
The timing of reafferent spikes from KO electroreceptors is shifted by EOD elongation through testosterone treatment. (A) Schematic representation of electrophysiological recording from KOs. Recording electrode is positioned over an individual KO without touching it. (B) Example traces in response to self-generated EODs recorded from an identical KO of a testosterone-treated fish at 1, 6, and 14 days after treatment. Inverted EOD waveform recorded from the same fish on the same day was used for sensory stimulation. Spike rate was calculated using a spike density function (see Materials and Methods). Raster plots show spike timing over 50 repetitions. (C) Daily changes in KO peak latency by vehicle (black) and testosterone (magenta) treatment. Each line connects points that correspond to the same KO recorded across multiple days. (D) Relationship between delay to EOD peak 1 and KO peak latency. Regression line (blue) was determined using a linear mixed-effect model. The slope is 1.11 and the intercept is 0.16. (E) Relationship between delay to EOD peak 1 and KO peak latency to EOD peak 1. The slope is 0.11 and the intercept is 0.16.

KOs typically emit a single spike with a fixed delay relative to EOD peak 1 (31, 32), but our recordings included several KOs that had a relatively strong second peak in their average firing rate (or spike density function; see Materials and Methods) (SI Appendix, Fig. S1A). We counted KOs with a second peak whose z-score was greater than 3 and whose timing was within 5 ms of EOD onset and found more testosterone-treated KOs (5/16 total KOs, 9/38 total responses over multiple days) in this range than vehicle-treated KOs (1/16 total KOs, 1/34 total responses) (SI Appendix, Fig. S1B). These second peaks resulted from either: (i) multiple spikes to a single EOD stimulus; or (ii) a single spike with variable timing across trials.

Overall, these results suggest that reafferent input to the KO sensory pathway can be altered by testosterone treatment, which may require coordinated changes to corollary discharge timing to inhibit it effectively.

### Testosterone shifts corollary discharge timing

To measure the inhibitory effect of corollary discharge in the KO pathway, we recorded evoked potentials from the anterior exterolateral nucleus (ELa) in the midbrain, which is the main target of nELL projection axons (Fig. 1B). In this preparation, neuromuscular paralysis blocks EOD production while spontaneous EOD commands remain, leaving the corollary discharge effects on sensory processing intact (24). We stimulated with 0.2 ms bipolar square electric pulses delivered with a specific delay, typically between 0.2 and 10.2 ms after the EOD command (EODC) recorded from spinal electromotor neurons (EMNs) (Fig. 4A). Electrosensory responses of a vehicle-treated fish were blocked for a narrow range of stimulus delays (∼3–4 ms) following the EOD command, due to the corollary discharge inhibition in nELL (Fig. 4B, left), as shown previously (31). Strikingly, the window over which responses were blocked in a testosterone-treated fish was delayed and elongated (Fig. 4B, right).

**Fig. 4.**
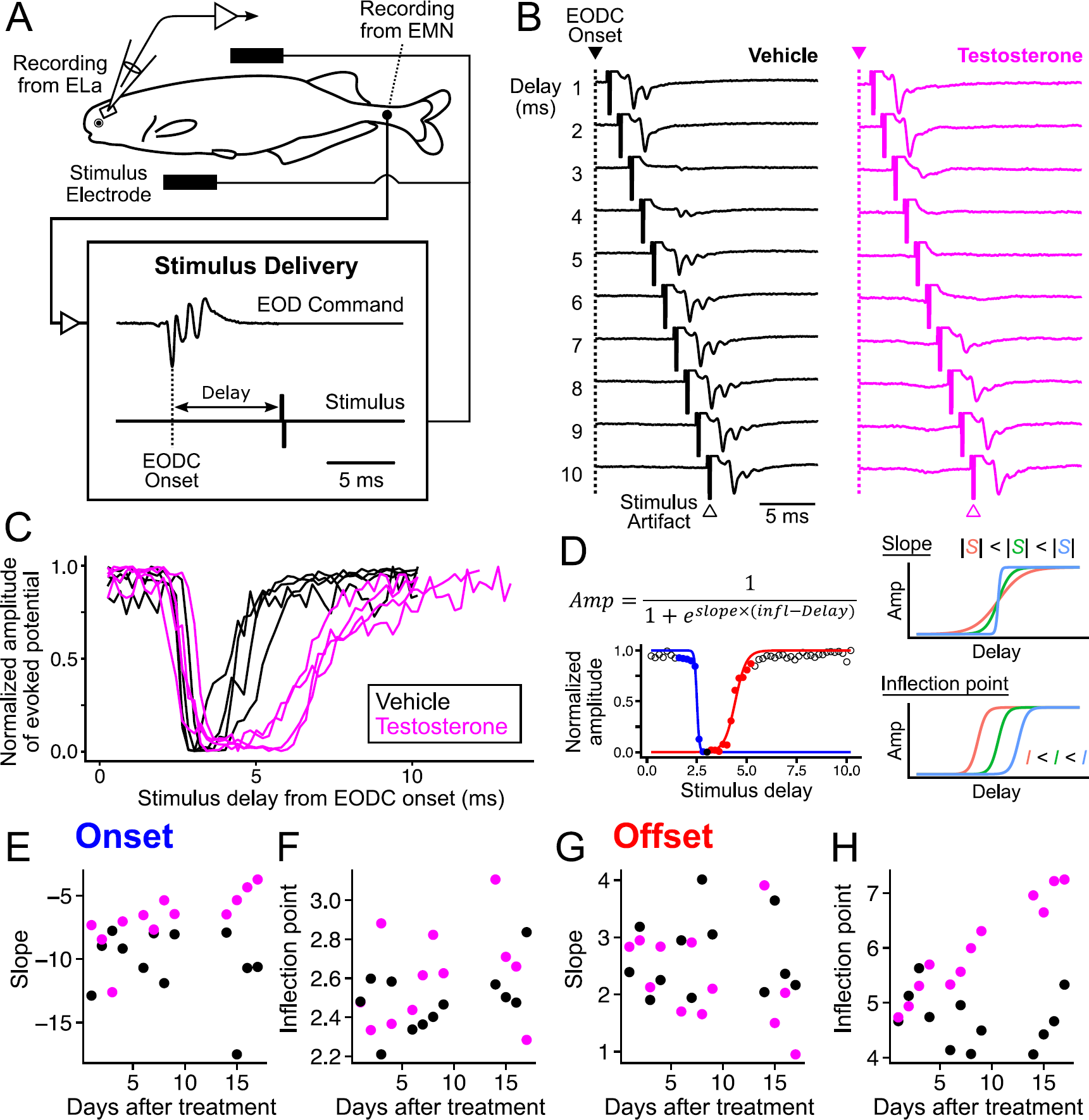
Corollary discharge timing is altered by testosterone treatment. (A) Measurement of corollary discharge timing. Although the fish is curarized to eliminate movement and silence EOD production, EOD commands (EODC) from spinal electromotor neurons (EMN) can be recorded as fictive EODs. Electrosensory stimuli can be delivered at fixed delays relative to the EODC onset, which is determined as the first negative peak, while recording evoked potentials in ELa. (B) Representative mean evoked potentials in response to stimuli at varying delays following the EOD command (1–10 ms) in vehicle- and testosterone-treated fish recorded 16 days after treatment. (C) Corollary discharge inhibition curves in vehicle- and testosterone-treated fish recorded 14–17 days after treatment. Normalized amplitudes were calculated using the maximum and minimum peak-to-peak amplitude across all stimulus delays. (D) Sigmoid curve fitting. On the left is the expression used and an example of an inhibition tuning curve fitted to separate onset and offset sigmoid curves. The filled black circle is the minimum point, with the left side defined as the onset direction and the right side as the offset direction. Filled colored circles are data points used for each sigmoid fit (blue, onset; red, offset). On the right is a description of how the coefficients, slope and inflection point, are related to the shapes of the curves. (E–H) Changes in onset slope (E), inflection point of the onset (F), offset slope (G), and inflection point of the offset (H).

From the evoked potential traces, we calculated the normalized amplitude across stimulus delays and generated an inhibition curve (Fig. 4C). We then divided the curve into onset and offset curves based on the point of minimum amplitude and fitted each curve to a sigmoid to determine two coefficients, slope and inflection point (Fig. 4D; see also Materials and Methods). We found that onset slope was flattened by testosterone (Fig. 4E; p = 0.0013 for treatment, p = 0.39 for days after treatment, p = 0.024 for the interaction, linear model [LM]), while the onset inflection point was not significantly affected (Fig. 4F; p = 0.15 for treatment, p = 0.15 for days after treatment, p = 0.89 for the interaction, LM). By contrast, the offset slope was not significantly affected by testosterone (Fig. 4G; p = 0.25 for treatment, p = 0.40 for days after treatment, p = 0.35 for the interaction, LM), but the offset inflection point was substantially delayed (Fig. 4H; p < 0.0001 for treatment, p = 0.0003 for days after treatment, p < 0.0001 for the interaction, LM).

### Shifted corollary discharge matches the shift in reafferent timing

We further examined whether the shifted corollary discharge was matched to the shifted reafferent input. Prior to evoked potential recording, we measured EOD timing relative to the EOD command in unparalyzed fish to compare the time courses of corollary discharge and EOD production (Fig. 5A) (31). We found no hormonally induced shift in EOD onset relative to the EOD command (Fig. 5B; p = 0.36 for treatment, p = 0.0077 for days after treatment, p = 0.73 for the interaction, LM). There was a slightly positive slope of onset versus day. The reason for this is unclear but may be due to the small amounts of ethanol used to dissolve the testosterone that were also provided to the vehicle treatment group. Importantly, the delay to EOD peak 1 from the EOD command was shifted by testosterone (Fig. 5C; p < 0.0001 for treatment, p = 0.0001 for days after treatment, p = 0.046 for the interaction, LM), demonstrating that EOD elongation indeed caused a shift in reafferent timing.

**Fig. 5.**
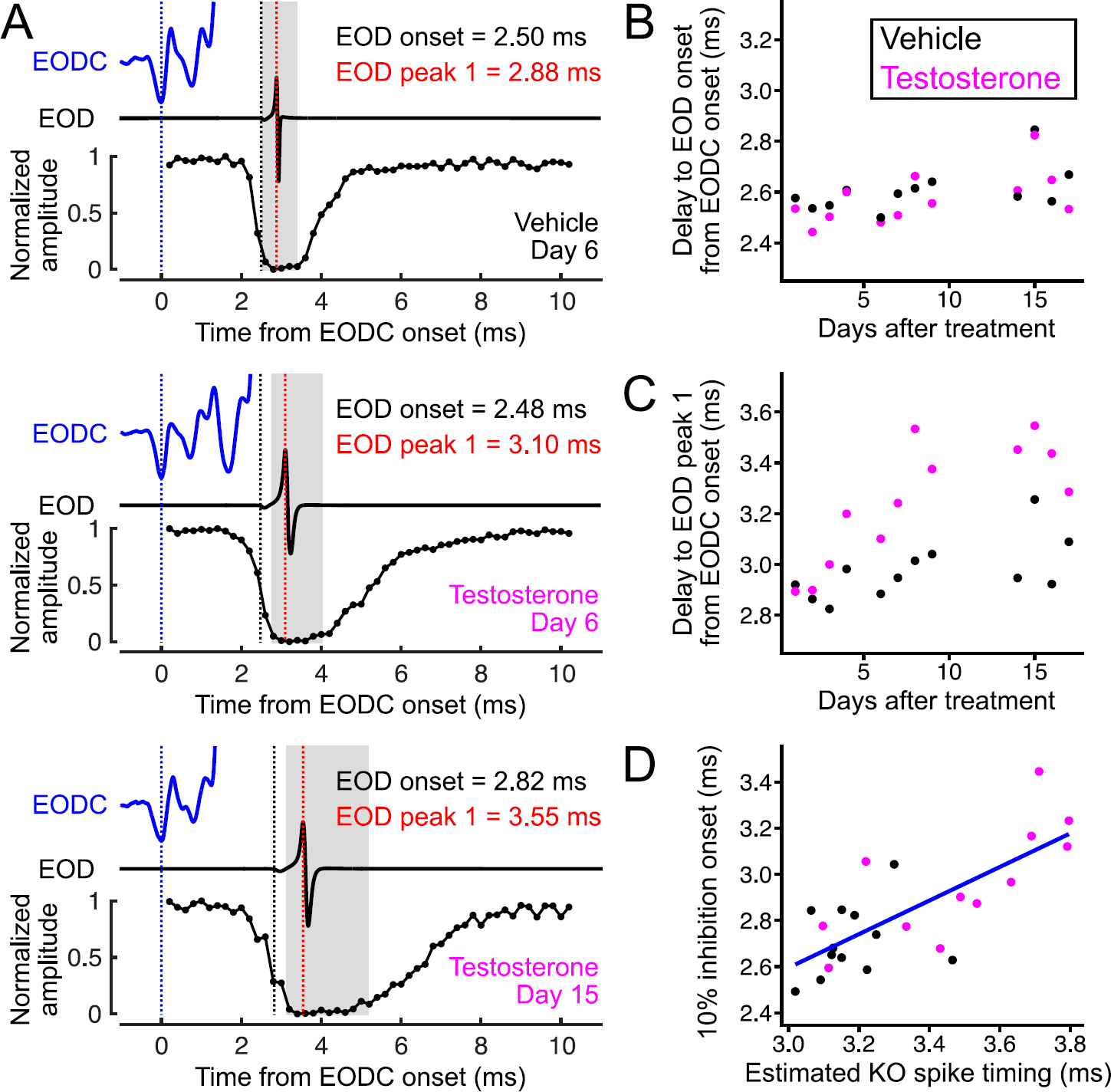
Shifted corollary discharge matches shifted reafferent spike timing. (A) Comparison of the time courses of the EOD command (EODC) (top blue traces), the EOD (middle trace), and corollary discharge inhibition. Vertical dotted lines indicate EODC onset (blue), EOD onset (black), and EOD peak 1 (red). Gray areas indicate 10% inhibition window calculated from the sigmoid curve fitting. (B) Daily change in EOD onset relative to EODC onset. (C) Daily change in EOD peak 1 relative to EODC onset. (D) Relationship between estimated KO spike timing and 10% inhibition onset. Regression line (blue) was determined using a linear model. The slope is 0.73 and the intercept is 0.41.

Given the KO recording data, we estimated KO reafferent spike timing for these fish (Fig. 3D, see also Materials and Methods). We found that the timing of strong inhibition onset, or 10% inhibition onset, tightly correlated with the estimated KO reafferent spike timing (Fig. 5D, R = 0.75, p < 0.0001). The slope and intercept of the regression line were 0.73 and 0.41, respectively.

### Sensory feedback is not necessary for corollary discharge shift

How is the match between EOD elongation and corollary discharge shift achieved? One possibility is that the altered sensory feedback tunes corollary discharge timing through plasticity or learning. To test this, we made fish electrically silent by spinal cord transection and measured corollary discharge timing by recording evoked potentials.

Because the transection also eliminated EOD commands from spinal EMNs, we recorded fictive EODs as field potentials from the command nucleus (CN) in the hindbrain (Fig. 6A), which fires one-to-one with EOD output in intact fish (33, 34) (Fig. 6B). CN activity was intact in all fish tested. Since CN potentials precede EOD commands from the EMN in intact fish, the apparent inhibition window by corollary discharge was delayed, but the differences between vehicle and testosterone fish were similar to those of intact fish (Fig. 6C, D).

**Fig. 6.**
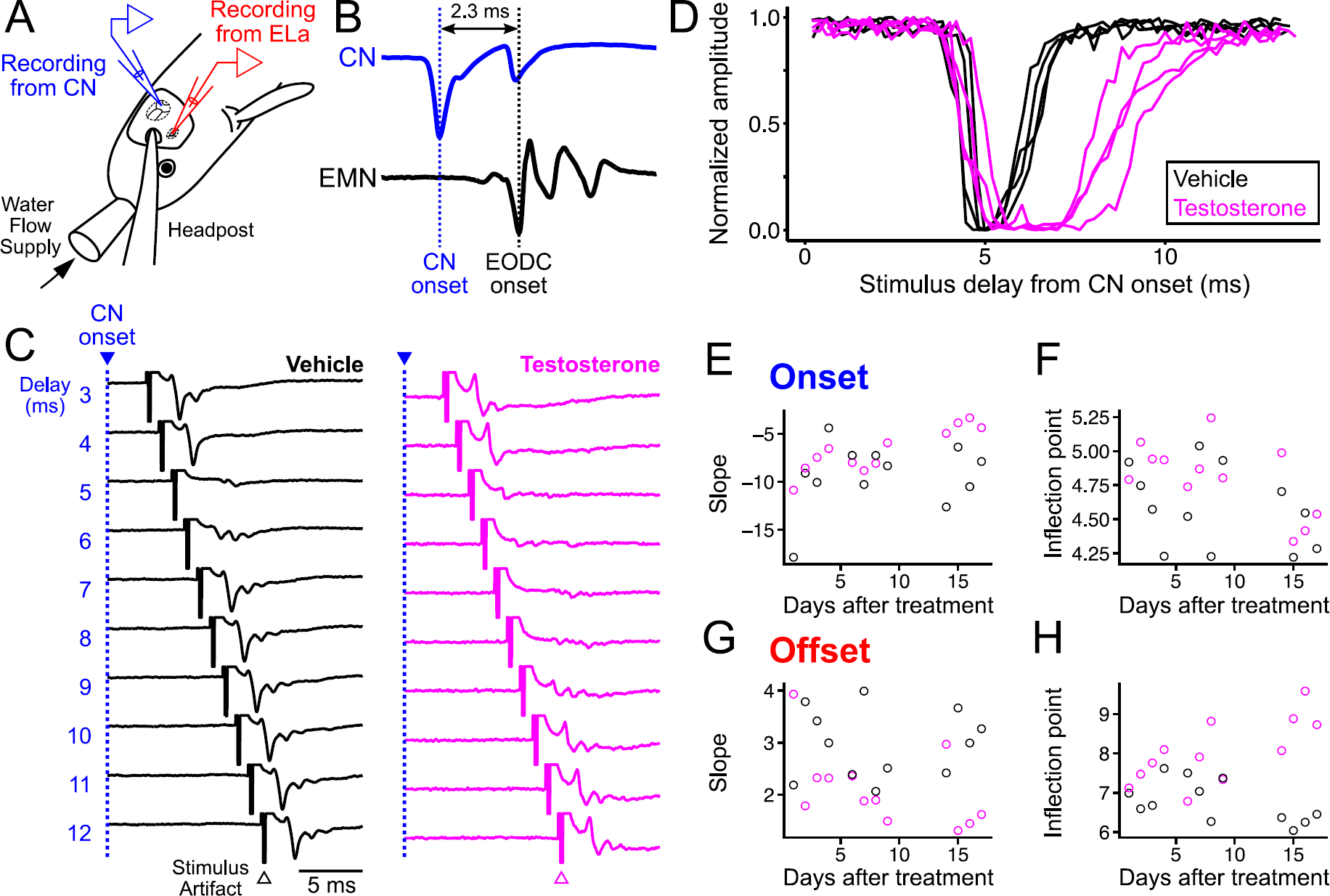
Testosterone alters corollary discharge without sensory feedback. (A) Measurement of corollary discharge timing from surgically silenced fish. Instead of EOD commands from EMNs, field potentials from the command nucleus (CN) in the hindbrain were used as a motor reference signal. (B) Example traces of a CN potential and EOD command from an intact fish. The CN potential typically has a double negative peak and its onset was determined as the first negative peak. EOD command signals from the EMNs follow CN onset with a fixed delay in intact fish. (C) Representative mean evoked potentials in response to stimuli at varying delays following the EOD command (3–12 ms) in vehicle- and testosterone-treated fish recorded 16 days after treatment. (D) Corollary discharge inhibition curves in vehicle- and testosterone-treated, surgically silenced fish recorded 14–17 days after treatment. (E–H) Changes in onset slope (E), inflection point of the onset (F), offset slope (G), and inflection point of the offset (H).

As in intact fish, we found that testosterone flattened the onset slope (Fig. 6E; p = 0.025 for treatment, p = 0.024 for days after treatment, p = 0.253 for the interaction, LM) and substantially delayed the offset inflection point (Fig. 6H; p < 0.0001 for treatment, p = 0.21 for days after treatment, p = 0.0008 for the interaction, LM). However, unlike intact fish, the interaction effect on onset slope was not significant. In addition, we found significant effects of testosterone on the onset inflection point (Fig. 6F; p = 0.043 for treatment, p = 0.023 for days after treatment, p = 0.60 for the interaction, LM) and on the offset slope (Fig. 6G; p = 0.0057 for treatment, p = 0.25 for days after treatment, p = 0.17 for the interaction, LM), which were also different from the results in intact fish.

To statistically test whether the effects of testosterone were influenced by sensory feedback, we made additional models that included both intact and silent fish, and added an additional independent variable, “silencing” (see also SI Appendix, Table S1). We found no significant interaction effects between treatment and silencing nor between treatment, silencing, and days (SI Appendix, Table S1), suggesting that absence of sensory feedback had little effects on the hormonal modulations. In these models, we found significant hormonal modulations including gradual changes in onset slope (p = 0.0001 for treatment, p = 0.017 for the interaction between treatment and days) and offset inflection point (p < 0.0001 for treatment, p < 0.0001 for the interaction), and relatively rapid changes in onset inflection point (p = 0.012 for treatment, p = 0.61 for the interaction) and offset slope (p = 0.0054 for treatment, p = 0.10 for the interaction). We also found a significant interaction effect between surgery and days on onset inflection point (p = 00070), suggesting that surgical silencing or an absence of sensory feedback might have slightly reduced the corollary discharge delay.

## Discussion

We found that testosterone modulates the timing of corollary discharge in a mormyrid fish (Fig. 4). This modulation corresponds to the hormonally induced change in EOD duration that shifts the reafferent spike timing of electroreceptors (Fig. 2, 3, and 5). Recordings from surgically silenced fish showed that exposure to sensory feedback is not necessary to drive this hormonal modulation of corollary discharge (Fig. 6). These results suggest that testosterone directly and independently adjusts the internal signal that predicts the sensory consequences of peripheral motor output.

Changes in circulating steroid hormone levels can modulate motor systems (2–6, 35), sensory systems (8–10), or both (36, 37), which is particularly evident in communication systems. These changes should alter the encoding of reafferent inputs while the nervous system still faces the challenge of distinguishing between self and other. Corollary discharges from motor centers serve as predictive signals of the timing of motor output and mediate this discrimination in sensory processing across modalities and species (24, 38–40). Mismatch between sensory prediction and actual sensory feedback is associated with hallucinations in patients with schizophrenia (41). Our results show that testosterone-treated mormyrid fish have modified a filter in their corollary discharge to match altered reafferent input (Fig. 5). The onset of this filter occurs just prior to the peak timing of the reafferent spikes, and its longer duration can cover more variable spike timings, which were more often observed in testosterone-treated KOs (Figs. 3, 4, and SI Appendix, Fig. S1). As the task of self-other discrimination is ubiquitous across species and sensory modalities, we expect that hormonal shifts in corollary discharge will be found in a wider range of systems (24, 40).

To minimize mismatch throughout sensorimotor circuits, coordination of hormonal effects on different circuit components is essential. For example, in a gymnotiform electric fish that emits highly regular EODs, androgen treatment increases EOD duration while simultaneously decreasing EOD frequency in a coordinated manner (35). Interestingly, localized androgen treatment of the electric organ increases EOD duration without affecting EOD frequency, suggesting independent regulation of EOD duration and frequency by androgens acting in the peripheral and central nervous systems, respectively, similar to our findings (42). In addition, androgens directly lower the frequency tuning of electroreceptors in gymnotiforms (36, 43), in contrast to the indirect hormonal effect on KO receptors in mormyrid fish (30).

We found that corollary discharge timing is regulated by circulating testosterone levels to achieve coordination of the electrocommunication circuit. Thus, one or more sites in the corollary discharge pathway may express androgen receptors. The corollary discharge originates in the CN and reaches the nELL via three additional nuclei, the bulbar command-associated nucleus (BCA), the mesencephalic command-associated nucleus (MCA), and the sublemniscal nucleus (slem) (Fig. 1B) (21, 23–25). A previous study examined androgen binding sites in the mormyrid brain but did not find binding sites in the corollary discharge pathway (44). However, the examined area was limited to the relatively ventral side and likely missed parts of this corollary discharge pathway (44). In addition, androgen binding sites may only become visible after testosterone treatment: a previous study using a gymnotiform fish showed that the expression of androgen receptors was upregulated following 15 days of hormone treatment (45). Moreover, the actions of testosterone may be mediated by aromatase, which converts testosterone into estrogen (46). Future studies should examine the distribution of both androgen and estrogen receptors throughout the corollary discharge pathway using testosterone-treated fish to identify where in the pathway they may be responsible for shifting corollary discharge timing.

The cellular mechanisms by which testosterone shifts corollary discharge timing remain to be determined. There are many possibilities. For example, testosterone, or estrogen produced by aromatase, may regulate GABA transmission in the synapses between slem and nELL, which could shape the time course of corollary discharge inhibition (47, 48). In addition, testosterone may change axonal morphology and myelination in the corollary discharge pathway to delay action potential propagation and the timing of inhibition (22, 49–51). Testosterone could also alter the passive or active electrical properties of neurons in this pathway to alter their excitability (7, 52, 53). A recent study documented changes in gene expression induced by testosterone in the electric organ of the same species of mormyrid we studied, many of which are likely associated with EOD elongation (54). It will be interesting to determine whether similar molecular cascades underlie shifts in corollary discharge timing.

Altered sensory feedback itself could provide a basis for adjusting corollary discharge through plasticity. Such plasticity has been found in several systems, including passive and active electrolocation systems in mormyrid fish (55–58). In these electrolocation systems, corollary discharge functions to subtract predictive sensory feedback from reafferent input by a modifiable efference copy, forming a “negative image” through cerebellum-like circuitry in the electrosensory lateral line lobe (24, 55, 56). This negative image is formed at synapses between parallel fibers that carry corollary discharge inputs from the CN and medial ganglion cells that receive sensory inputs through spike-timing-dependent plasticity (59–62). By contrast, in the communication pathway, we found that removal of sensory feedback had no effect on hormonal modulations of corollary discharge (Fig. 4 and 6). However, we cannot conclude that sensory feedback has no effect on corollary discharge inhibition because we did not test whether altered sensory feedback is sufficient to modify corollary discharge inhibition. We observed very similar inhibition curves in intact and silent fish (Fig. 4 and 6), but it is also possible that altered sensory feedback plays a key role in fine-tuning the onset of inhibition, which was fixed to KO spike timing in response to simulated reafferent input in intact fish (Fig. 5). More localized treatment of testosterone in the electric organ, or other manipulations that alter sensory feedback over long periods of time without increasing systemic levels of testosterone, may allow us to test the effects of altered sensory feedback.

Here we demonstrate hormonal regulation of corollary discharge in concert with EOD elongation. Coordinated changes in corollary discharge are essential for adaptive behavioral change not only through hormonal plasticity, but also through development or evolution, because all animals are constantly faced with the task of discriminating between self and others (24, 38–40). Coordination of behavioral change and a corollary discharge shift is also found in the diversification of communication signals among mormyrid species, and in developmental changes in these signals within species (31, 63). It will be very interesting to test whether these evolutionary and developmental shifts, and the hormonal shifts we describe here, share the same substrates or mechanisms responsible for the precise matching of corollary discharge and reafferent timing. Mormyrids are an excellent system to study sensorimotor coordination underlying plastic behavior and its relationship to evolutionary change.

## Methods

All procedures were in accordance with guidelines established by the National Institutes of Health and were approved by the Animal Care and Use Committee at Washington University in St. Louis.

### Animals

We used a total of 54 *Brienomyrus brachyistius* of both sexes in non-reproductive state (5.4–10.9 cm in standard length). All fish were purchased from Bailey Wholesale Tropical Fish or Alikhan Tropical Fish. The fish were housed in groups with a 12 h: 12 h light/dark cycle, temperature of 25–29°C, pH of 6–7, and water conductivity of 200–400 µS/cm. Fish were fed live black worms or frozen blood worms four times per week.

Half of the fish were testosterone treated and the other half were vehicle treated. All experiments below used the same number of fish for each treatment. Six fish were used for KO recording. Another twenty-four fish were used for simultaneous recording of EOD and EODC and evoked potential recording from ELa. For 8 of these fish, the change in EOD was followed up to 13 days after the start of treatment. The remaining 24 fish were surgically silenced to eliminate EOD production by spinal cord transection before treatment, and then used for evoked potential recording.

### Surgery for silencing the EOD

Fish were anesthetized in a solution of 300 mg/L MS-222 (Sigma Millipore). When movement ceased, the fish were removed from the solution and 70% ethanol was applied around the incision site, just anterior to the electric organ. A 26-gauge needle was then inserted through the skin to transect the spinal cord. After transection, lidocaine (2% solution; Radix Laboratories) was applied around the incision site, but not directly at the incision site, for local anesthesia. Finally, the incision was closed with a small amount of superglue and the fish was returned to its home tank. This operation did not affect motility, as the spinal motorneurons controlling movement are all located anterior to the incision site (30, 64). Prior to treatment, surgically silenced fish were allowed to recover in isolation for at least 6 days.

### Hormone treatment

A treatment tank (25.4 * 30.5 * 50.8 cm) was prepared for each treatment group, divided into four compartments with mesh panels, and filled with 30 L of treated aquarium water. For testosterone fish, 60 mg of solid 17α-methyltestosterone (Sigma-Aldrich) dissolved in 0.4 mL 95% ethanol was added to the testosterone tank on the start day and the next day, and then every two days thereafter. For vehicle fish, only 0.4 mL 95% ethanol was added to the vehicle tank on the same schedule as the testosterone treatment. Except for the treatment and separation with mesh panels, tank conditions were identical to our normal fish housing.

### EOD recording

EOD recordings were made from freely swimming fish prior to EOD and EODC recording or KO recording in intact fish. EODs were amplified 10 times, bandpass filtered (1 Hz–50 kHz) (BMA-200, CWE), digitized at a rate of 195 kHz (RP2.1, Tucker-Davis Technologies), and stored using custom software in MATLAB (The MathWorks). We measured EOD duration and delay to peak 1 according to previously defined criteria (31). Peak power frequency was calculated by fast Fourier transformation. Each value was calculated from 10 EODs obtained from each fish, and the average was used for analysis.

### KO recording

The recording method was similar to previous studies (65, 66). Fish were anesthetized with a solution of 300 mg/L MS-222, and then paralyzed and electrically silenced with 50–60 µL of 0.05 mg/mL gallamine triethiodide (Flaxedil, Sigma-Aldrich). The fish were then placed on a plastic platform with lateral supports in a recording chamber (20 * 12.5 * 45 cm) filled with freshwater covering the entire body of the fish and were respirated with aerated freshwater through a pipette tip placed in the fish’s mouth. To verify that the fish had recovered from anesthesia, field potentials were recorded from EMNs using a pair of electrodes placed next to the fish’s tail. After recovery, indicated by EOD commands from the EMNs, the recording session was started. After recording, the fish were allowed to fully recover from paralysis before being returned to their home tank.

We made recording electrodes from borosilicate capillary glass (o.d. = 1 mm, i.d. = 0.5 mm; Model 626000, A-M Systems). Using a Bunsen burner, we bent the last ∼1–2 cm to a ∼10–30-degree angle and polished the tip. The electrode was filled with tank water, placed in an electrode holder with a Ag-AgCl wire connected to the headstage of an amplifier (Neuroprove Model 1600, A-M Systems) and positioned over individual KOs without touching them. The extracellular activity was referenced to ground, amplified 10 times and low-pass filtered (cut-off frequency = 10 kHz) with the amplifier, and digitized at a sampling rate of 97.7 kHz (RP2.1, Tucker-Davis Technologies). Electrosensory stimuli were generated at a sampling rate of 195.31 kHz (RP2.1, Tucker-Davis Technologies), attenuated (PA5, Tucker-Davis Technologies), and delivered through the recording electrode as constant-current stimuli. We used a bridge balance to minimize stimulus artifact. Recording traces were stored using custom MATLAB (The MathWorks) software from 20 ms before stimulation to 20 ms after stimulation.

We used an inverted (or head-negative) EOD waveform for stimulation, which was recorded from the same individual just prior to recording. When a fish generates an EOD, all the KOs on its skin receive the same-direction currents consisting of a large outward current followed by a large inward current. This waveform is opposite to the waveform obtained when a recording electrode is placed at the head and a reference electrode is placed at the tail. We tested each KO with at least two different stimulus intensities, with a 5 dB attenuation interval, including one intensity where KO spike amplitude exceeded the stimulus artifact and one where it did not. We presented 50 repetitions for each stimulus intensity.

For each recording, we selected the maximum stimulus intensity at which spikes could be reliably distinguished from artifact for further analysis. Spikes were detected by finding the peak voltage that crossed a manually set threshold specific to each KO. For some KOs, a stimulus artifact template was created by taking the median trace of non-responsive traces to weaker stimuli and scaling it. The template trace was subtracted from the recording traces to stronger stimuli and spikes were detected by the threshold crossing method.

We computed the spike density function (SDF) by convolving each KO spike time with a Gaussian of 0.1 ms width and then averaging over stimulus repetitions. We first determined the timing of the largest peak for each KO SDF trace as the first peak. KO peak latency was calculated as the interval between EOD stimulus onset and the time of the first peak. KO peak latency to EOD peak 1 was calculated as the interval between the time of EOD peak 1 and the time of the first peak.

When looking at the distribution of KO peak latency to EOD peak 1, two obvious outliers were found. These KO recordings showed weak responses to the stimuli and were excluded. As a result, a total of 72 recordings from 32 different KOs were used for the subsequent statistical analyses. Note that the 23 out of 32 KOs could be recorded on multiple days.

We also determined the timing of the second largest peak for each SDF trace as the second peak. The z-score of the second peak was calculated using the z-score over each SDF.

### EOD and EOD command recording

The recording and analysis methods were similar to a previous study (31). Fish were anesthetized in a solution of 300 mg/L MS-222 (Sigma Millipore) and placed on a plastic platform with lateral supports in the recording chamber filled with freshwater covering the entire body of the fish. Fish were restrained by lateral plastic pins, a plastic tube on the tail, and two or three folded paper towels on the dorsal skin surface. EOD commands from spinal EMNs were recorded with a pair of electrodes located within the plastic tube and oriented parallel to the fish’s electric organ, amplified 1000×, and bandpass filtered (10 Hz to 5 kHz) (Model 1700, A-M Systems). While EOD commands from EMNs were recorded, the EODs were recorded by separate electrodes, amplified 10 times, and bandpass filtered (1 Hz to 50 kHz) (BMA-200, CWE). These recordings were digitized at a rate of 1 MHz and saved (TDS 3014C, Tektronix).

EOD command traces from EMNs were averaged across trials, and EOD traces were filtered by a 21st-order median filter whose time window was 0.02 ms and averaged across trials. EOD onset was determined in the same way we determined EOD onset in freely swimming EOD recordings. EOD command onset was determined as the first negative peak in the averaged EMN trace. Delay to EOD onset was calculated as the time between EOD command onset and EOD onset. Delay to EOD peak 1 from the EOD command was calculated as the sum of the delay to EOD onset and the delay between EOD onset and peak 1 recorded from freely swimming fish.

### Evoked potential recording from intact fish

The recording and analysis methods were similar to a previous study (20, 22, 31). Fish were anesthetized with a solution of 300 mg/L MS-222, and then paralyzed with 0.05–0.1 mL of 3.0 mg/mL Flaxedil. The fish were then transferred to a recording chamber filled with water and positioned on a plastic platform, leaving a small region of the head above water level. During surgery, we maintained general anesthesia by respirating the fish with an aerated solution of 100 mg/ml MS-222 through a pipette tip placed in the mouth. For local anesthesia, we applied 0.4% lidocaine on the skin overlying the incision site, and then made an incision to uncover the skull overlying the ELa. Next, we glued a headpost to the skull before using a dental drill and forceps to remove a rectangular piece of skull covering the ELa. After exposing ELa, we placed a reference electrode on the nearby cerebellum. Following surgery, we switched respiration to fresh water and allowed the fish to recover from general anesthesia. EOD commands were also recorded from the EMNs and sent to a window discriminator for time stamping (SYS-121, World Precision Instruments). At the end of the recording session, the respiration of the fish was switched back to 100 mg/L MS-222 until no EOD commands could be recorded, and then the fish was euthanized by freezing.

Recording electrodes (Model 626000, A-M Systems) were pulled with a micropipette puller (Model P-97, Sutter Instrument), broken to a tip diameter of 10–20 µm, and filled with 3 M NaCl solution. Evoked field potentials were amplified 1000×, bandpass filtered (10 Hz–5 kHz) (Model 1700, A-M Systems), digitized at a rate of 97.7 kHz (RX8, Tucker-Davis Technologies), and stored using custom software in MATLAB (The MathWorks). We presented 0.2 ms bipolar square pulses at several delays following the EOD command onset, typically ∼0.2–10.2 ms with 0.2 ms intervals, in a randomized order. The stimulus intensity was fixed at 73.6 mV/cm. Each delay was repeated 10 times.

Technically, the window discriminator could not detect the minimum point of the first negative peak in the EOD command, but rather the point at which a manually selected threshold was crossed. Therefore, we also recorded the EOD command and the digital output of the window discriminator to measure the stimulus delay to EOD command onset. Using this, the corrected stimulus delay was used in the subsequent analyses.

To characterize corollary discharge inhibition, we first calculated the normalized amplitude by the following steps: (i) calculating the peak-to-peak (PP) amplitude 2–4 ms after stimulus onset for each stimulus delay, (ii) subtracting the minimum PP amplitude across all delays, and (iii) dividing by the difference between the maximum PP amplitude and the minimum PP amplitude. Sigmoid curve fitting was then applied to the normalized amplitude curve as follows: (i) determining the minimum point and dividing into onset and offset directions, (ii) selecting the points ranging from the minimum amplitude point to a point above 50% amplitude where the next point is lower than that point for the first time for each direction, and (iii) fitting the following expression:

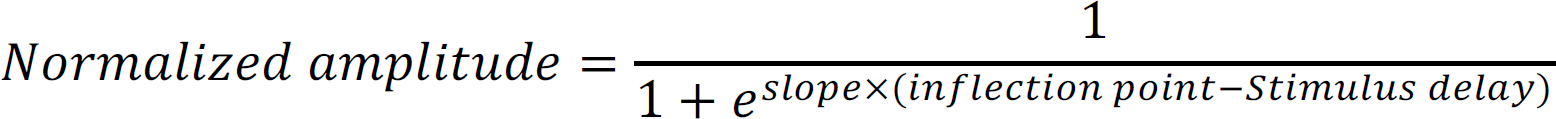

where slope and inflection point are the coefficients of this curve. The coefficients of the onset curve were used to calculate 10% onset and offset.

### Evoked potential recording from silent fish

The procedure was almost the same as in intact fish, but EOD commands from the EMNs were silent because the spinal cord had been cut anterior to the electric organ. As an alternative motor reference signal, we recorded field potentials from the command nucleus (CN) in the hindbrain. Before exposing ELa, we made a second incision window in the skull on the dorsal surface above the CN. After recovery from anesthesia, we searched for a CN potential using another electrode controlled by a motorized micromanipulator (MP-285, Sutter Instrument). CN typically exhibited a double negative potential, and the onset was determined as the first negative peak. Field potential signals from the CN were processed in the same way as EOD command recording from the EMNs to trigger electrosensory stimuli. Since the CN onset preceded the EMN onset, a wider range of stimulus delay to the command (∼0.2–13.2 ms with 0.2 ms intervals) was used. The stimulus delay from the CN field potential was also measured and the corrected stimulus delay was used in the analyses.

Our recordings of command-related potentials from the hindbrain possibly include the medullary relay nucleus that is immediately dorsal to the CN. Technically, it was difficult to accurately distinguish between these nuclei without the EMN reference. Because these nuclei are electrically coupled and have a very small difference in peak timing relative to the EMN in intact fish (33, 34), we analyzed the recordings without distinguishing between them.

### Statistical analyses

We performed all statistical tests using R version 4.0.3. We used the nlme package to make a model that accounts for random effects. To assess hormonal effects on EOD duration, peak power frequency and delay to EOD peak 1, we made linear mixed models in which the fixed effects were treatment (vehicle or testosterone), days after treatment, and the interaction, and the random effect was individual fish (Fig. 2B–D). To assess the hormonal effects on KO peak latency, we made a linear mixed model in which the fixed effects were treatment, days after treatment, and the interaction, and the random effect was individual KO (Fig. 3C). To assess the hormonal effects on the corollary discharge inhibition curves (Fig. 4 and Fig. 6) and EOD timing relative to the EOD command onset (Fig. 5B, C), we made linear models with fixed effects of treatment, days after treatment, and the interaction. Additionally, we also made linear models with fixed effects of treatment, surgery, days after treatment, and the interactions (Appendix SI, Table S1). To test for significant effects, we applied an analysis of variance to these models. To describe the relationship between delay to EOD peak 1 and KO peak latency (Fig. 3D) or KO peak latency to EOD peak 1 (Fig. 3E), we made a linear mixed model in which the fixed effect was delay to EOD peak 1 and the random effect was individual KO and determined the slope and intercept of the regression line. In this case, we tested whether a given parameter was significantly different from zero. Using the coefficients from Fig. 3D, we estimated KO spike timing of fish (intact) that were used for evoked potential recording (Fig. 5D). To determine the relationship between estimated KO spike timing, we calculated the Pearson’s correlation coefficient.

**Fig. S1.**
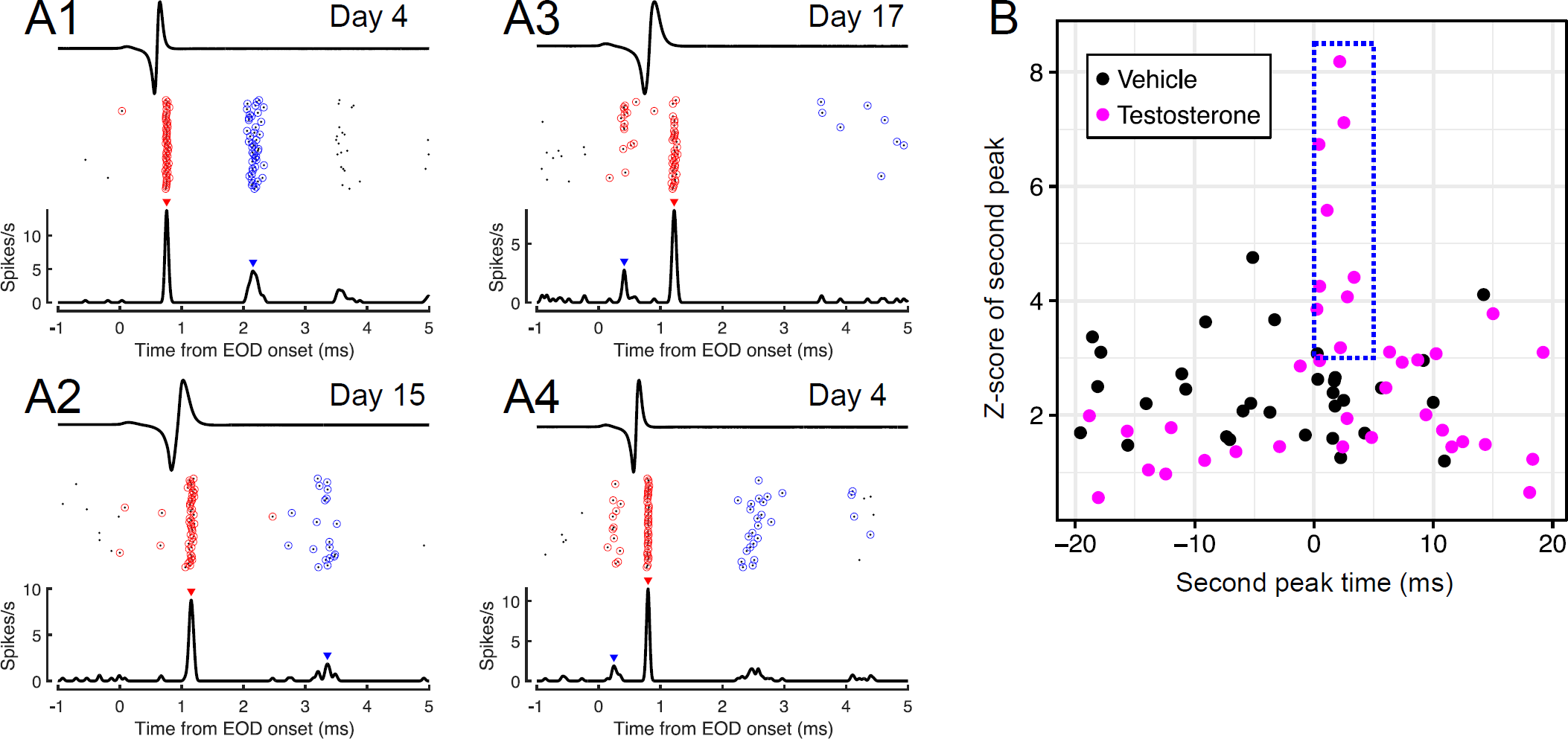
Several testosterone-treated KOs have multiple spike timings. (A) Four example traces. Upper traces show the electrosensory stimulus mimicking self-generated EODs similar to Fig. 2B. Middle raster plots show spike timing over 50 repetitions. Red circles indicate the first spike in each repetition and blue circles indicate the second spike after each EOD stimulus onset. Lower traces show spike rate averaged across repetitions. The red triangle above each rate trace indicates the maximum peak of KO response, and the blue triangle indicates the second maximum peak. The KOs in A1 and A2 frequently produced more than one spike in response to a single EOD stimulus. The KOs in A3 and A4 sometimes produced a spike following peak 0 of the EOD, seen as a small peak before peak 1, but did not respond to both peak 0 and peak 1. The KO in A4 sometimes produced an additional spike ∼2–3 ms after EOD stimulus onset similar to KOs in A1 and A2. (B) Scatter plot of z-score of second peak versus second peak time relative to EOD stimulus onset. The blue rectangle indicates a range in which the z-score is greater than 3 and the peak time is within 5 ms of the EOD stimulus onset. Note that three KOs (two vehicle-treated and one testosterone-treated) were excluded from this plot because they showed no spontaneous activity and produced a spike only following peak 1 of the EOD, making it impossible to detect a second peak.

**Table S1.** Analysis-of-variance table for linear models that test the significance of treatment, day, silencing and the interactions on parameters of corollary discharge inhibition curve. Single, double, and triple asterisks represent p < 0.05, p < 0.01, and p < 0.001, respectively.

| Response variable | Independent variable | Df | Sum Sq | Mean Sq | F value | Pr(>F) | Significance |
| --- | --- | --- | --- | --- | --- | --- | --- |
| Onset slope | Treatment | 1 | 113.9637475 | 113.9637475 | 18.44490306 | 0.000108325 | *** |
|  | Day | 1 | 35.90932305 | 35.90932305 | 5.811883141 | 0.020604649 | * |
|  | Surgery | 1 | 3.437660044 | 3.437660044 | 0.556381373 | 0.460083226 |  |
|  | Treatment:Day | 1 | 38.4517822 | 38.4517822 | 6.223377265 | 0.016837112 | * |
|  | Treatment:Silencing | 1 | 2.832352111 | 2.832352111 | 0.458412971 | 0.502263745 |  |
|  | Day:Surgery | 1 | 9.536640878 | 9.536640878 | 1.543494492 | 0.221332922 |  |
|  | Treatment:Day:Silencing | 1 | 3.309198257 | 3.309198257 | 0.535589978 | 0.468532447 |  |
|  | Residuals | 40 | 247.1441506 | 6.178603764 | NA | NA |  |
| Onset inflection point | Treatment | 1 | 0.371669027 | 0.371669027 | 6.859448749 | 0.012395966 | * |
|  | Day | 1 | 0.053436298 | 0.053436298 | 0.986209568 | 0.326640296 |  |
|  | Surgery | 1 | 55.19301677 | 55.19301677 | 1018.631208 | 4.49E-30 | *** |
|  | Treatment:Day | 1 | 0.013703654 | 0.013703654 | 0.252911875 | 0.617790099 |  |
|  | Treatment:Silencing | 1 | 0.031524124 | 0.031524124 | 0.581802888 | 0.450081542 |  |
|  | Day:Surgery | 1 | 0.437274797 | 0.437274797 | 8.070255634 | 0.007044994 | ** |
|  | Treatment:Day:Silencing | 1 | 0.006091576 | 0.006091576 | 0.112424895 | 0.739152344 |  |
|  | Residuals | 40 | 2.167340498 | 0.054183512 | NA | NA |  |
| Offset slope | Treatment | 1 | 4.522947748 | 4.522947748 | 8.621891587 | 0.00548433 | ** |
|  | Day | 1 | 1.065722574 | 1.065722574 | 2.031538944 | 0.161821566 |  |
|  | Surgery | 1 | 0.05814907 | 0.05814907 | 0.110846953 | 0.740920723 |  |
|  | Treatment:Day | 1 | 1.46400022 | 1.46400022 | 2.790757683 | 0.102617239 |  |
|  | Treatment:Silencing | 1 | 0.735551111 | 0.735551111 | 1.402147955 | 0.243353147 |  |
|  | Day:Surgery | 1 | 0.009775863 | 0.009775863 | 0.018635287 | 0.892101997 |  |
|  | Treatment:Day:Silencing | 1 | 0.02810665 | 0.02810665 | 0.05357844 | 0.818129586 |  |
|  | Residuals | 40 | 20.98355194 | 0.524588798 | NA | NA |  |
| Offset inflection point | Treatment | 1 | 20.06573828 | 20.06573828 | 91.33509195 | 7.02E-12 | *** |
|  | Day | 1 | 2.938131566 | 2.938131566 | 13.37376741 | 0.000735331 | *** |
|  | Surgery | 1 | 51.04867527 | 51.04867527 | 232.3630152 | 2.95E-18 | *** |
|  | Treatment:Day | 1 | 9.459451591 | 9.459451591 | 43.05746784 | 7.68E-08 | *** |
|  | Treatment:Silencing | 1 | 0.00149307 | 0.00149307 | 0.006796146 | 0.934708989 |  |
|  | Day:Surgery | 1 | 0.541616887 | 0.541616887 | 2.465328089 | 0.124261121 |  |
|  | Treatment:Day:Silencing | 1 | 0.00869524 | 0.00869524 | 0.039578933 | 0.843313925 |  |
|  | Residuals | 40 | 8.78774537 | 0.219693634 | NA | NA |  |

## Acknowledgement

This work was supported by the National Science Foundation (IOS-1755071 and IOS-2203122 to B.A.C.) and a Japan Society for the Promotion of Science Overseas Research Fellowship (202060318 to M.F.). We thank Martin W. Jarzyna for providing feedback on earlier versions of the manuscript and collecting data for Figure 6B, and Mauricio Losilla for sharing the hormone treatment protocol.

## Notes

### Competing Interest Statement

The authors have declared no competing interest.

