## Supplementary figures and images for "Hormonal coordination of peripheral motor output and corollary discharge in a communication system"

### Figure S1

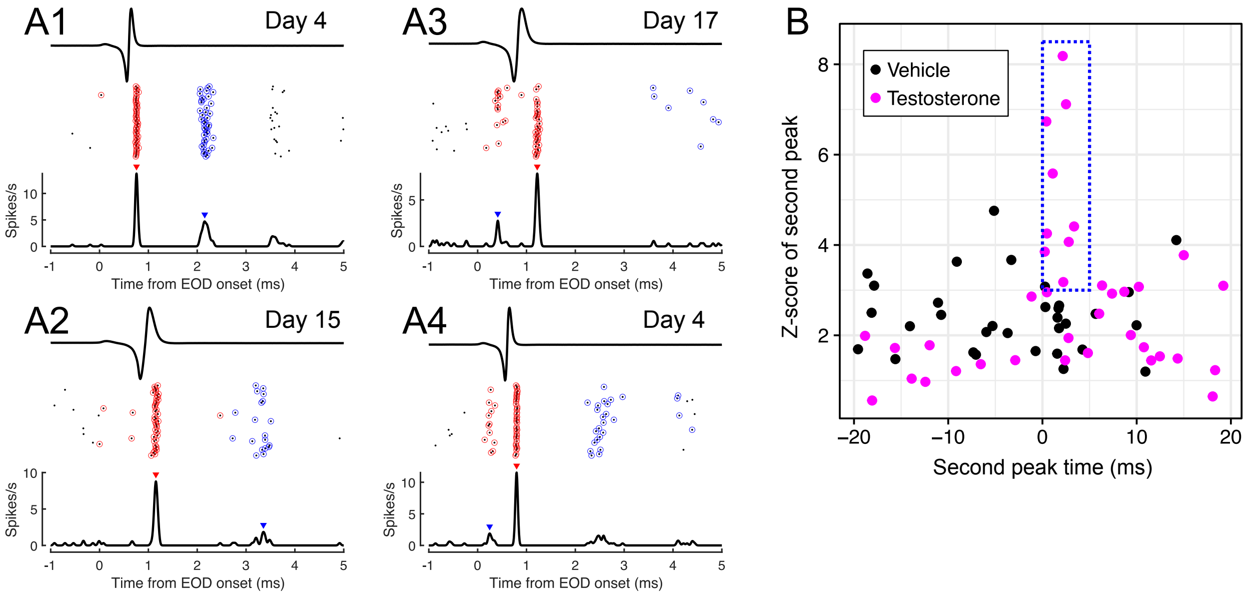
